# Changes of gene expression but not cytosine methylation are associated with plasticity of male parental care reflecting behavioural state, social context, and individual flexibility

**DOI:** 10.1101/139634

**Authors:** CB Cunningham, L Ji, EC McKinney, KM Benowitz, RJ Schmitz, AJ Moore

**Author notes:** Corresponding Author: C. B. Cunningham.

## Abstract

Behaviour is often on the front line of plasticity in response to different environments. At the genetic level, behavioural changes are likely to be associated with changes of gene expression. Most studies to date have focused on gene expression differences associated with discrete behavioural states reflecting development or age-related changes, such as honey bee castes. However, more rapidly flexible behaviour is often observed in response to social context or simple individual variation. The differences in genetic influences for the different forms of plasticity are poorly understood. In this study we contrasted gene expression during male parental care of the burying beetle, *Nicrophorus vespilloides*, in a factorial design. Male *N. vespilloides* males typically do not provide care when females are present. However, male care is inducible by the removing female and has parental effects equivalent to female care. We used this experimental manipulation to isolate gene expression and cytosine methylation associated with differences of behavioural state, differences of social context, or differences of individual flexibility for expressing care. The greatest number of differentially expressed genes was associated with behavioural state, followed by differences of social contexts, and lastly differences of individual variation. DNA methylation has been hypothesized to regulate the transcriptional architecture that regulates behavioural transitions. We tested this hypothesis by quantifying differences of cytosine methylation that were associated with differences of behavioural state and individual flexibility. Changes of cytosine methylation were not associated with changes of gene expression. Our results suggest a hierarchical association between gene expression and the different sources of variation that influence behaviour, but that this process is not controlled by DNA methylation despite reflecting levels of plasticity in behaviour. Our results further suggest that the extent that a behaviour is transient plays an underappreciated role in determining the molecular mechanisms that underpin the behaviour.

## Introduction

Behaviour, like all phenotypes, is traceable to how and when genes are expressed. Transcriptional profiling has revealed distinct transcriptional architectures associated with distinct behavioural states (Zayed and Robinson, 2012; Cardoso et al., 2015; Parker et al., 2015; Palmer et al., 2016; Jacobs et al., 2016), which is further reflected in protein abundance (Cunningham et al., 2017). However, the variation seen within and across behavioural states is not only a result of the gene expression that underpins the behaviour itself, but also reflects environmental factors, such as social context and individual variation in response to similar stimuli. An outstanding question is how much gene expression belongs to plasticity of behavioural state, social context, and individual flexibility. This question requires examining factors of interest at the same time and with experimental designs that minimize other differences that can exist when examining highly distinct behavioural states (Benowitz et al., 2017). Furthermore, the mechanisms that regulate gene expression variation are not fully characterised, but are needed for a full understanding of the evolution and mechanistic basis of behaviour.

In this study, we sought to partition the gene expression associated with three forms of behavioural plasticity: (1) the gene expression that reflects differences in behavioural state, (2) the gene expression that reflects response to social contexts, and (3) the gene expression that reflects individual behavioural variation (individual flexibility). We can examine all three of these factors simultaneously and partition their influences by manipulating male parental care behaviour of the subsocial beetle *Nicrophorus vespilloides*. This social behaviour displays considerable plasticity, making it productive for the investigation of transcriptional architecture of flexible social behaviours under different conditions. In this species, males but not female parental care is naturally plastic (Smiseth et al., 2005). With a mate, males can but do not always participate in the feeding of the offspring but instead provide indirect forms of care, such as excretion of anti-microbial compounds to cover the carcass (Smiseth et al., 2005). With the removal of his female mate, some males begin feeding offspring (Smiseth et al., 2005). Because we can manipulate male parental care *via* changes of social context, we can generate factorial crosses of males with or without mates that do or do not feed offspring. This helps us directly isolate potential effects on gene expression from behavioural state, social context, and individual flexibility displayed for those behaviours (Table 1). This is important because social behaviour is multifaceted and moving to factorial designs helps us to begin disentangling the influence of single variables, rather than comparing gene expression across behavioural states that necessarily differ for many variables (Lockett et al., 2012; Benowitz et al., 2017). We also have a sequenced and annotated genome for *N. vespilloides*, and there is gene body cytosine methylation (Cunningham et al., 2015).

**Table 1.**
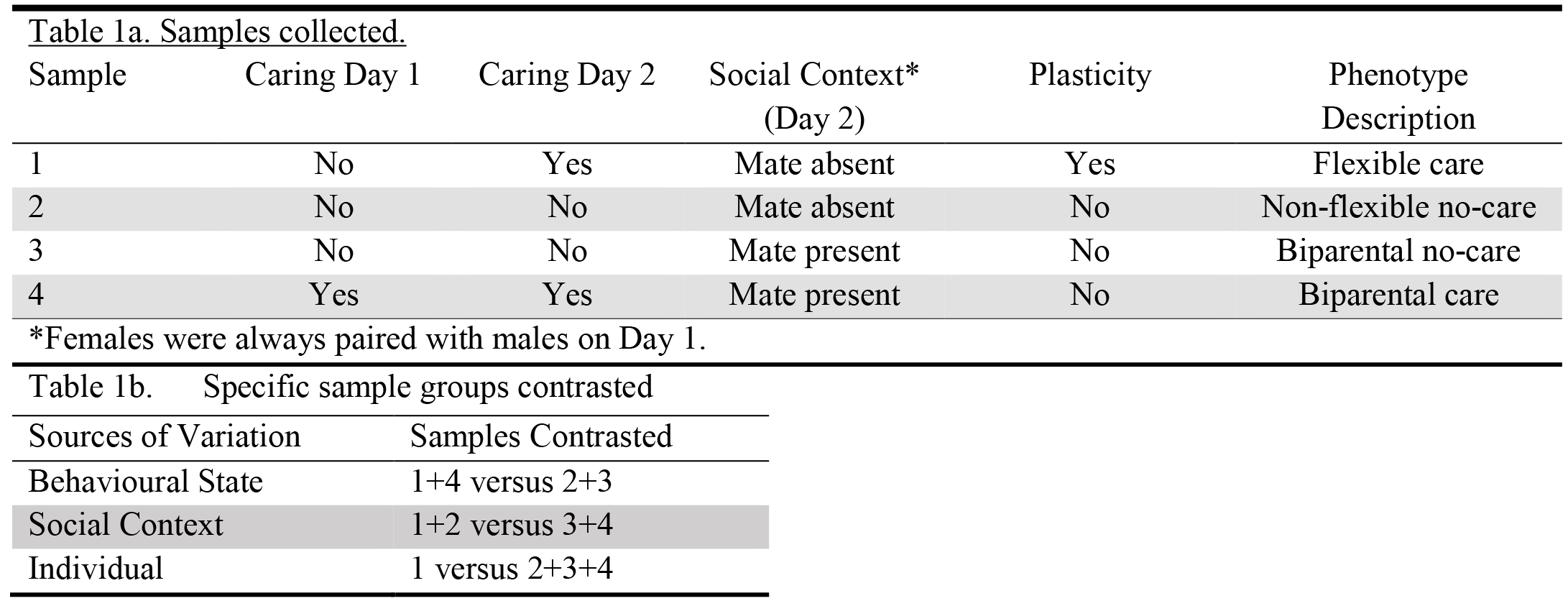
Experimental design. Four different sample groups collected, reflecting differences in male social context (presence or absence of the female parent) or parental behaviour (expressed or not expressed) on two days of observation. From these, three different contrasts were made.

Within this experimental design we also sought to test if DNA methylation regulated any rapid changes of gene expression during socially responsive parental care. DNA methylation is a core mechanisms regulating gene expression (Cardoso et al., 2015). It is stable (Turecki, 2014; Yan et., 2015), reversible (Yan et al., 2015), and can have relatively short-term turnover in animals (Levenson et al., 2006; Guo et al., 2011; Herb et al., 2012; Mizuno et al., 2012; Baker-Andresen et al., 2013; *contrario sensu* Cardoso et al., 2015). Therefore, cytosine methylation has excellent characteristics to regulate the gene expression underlying behaviours (Cardoso et al., 2015; Yan et al., 2015). Cytosine methylation is associated with different behaviours of a range of insects, including different hymenopterans (Kucharski et al., 2008; Lyko et al., 2010; Bonasio et al., 2012; Foret et al., 2012; Herb et al., 2012; Lockett et al., 2012; Amarasinghe et al., 2014; Kucharski et al., 2016) and an orthopteran (Wang et al., 2014). However, its association with behaviour is not ubiquitous, as cytosine methylation is not associated with different behaviours of several hymenopterans (Patalano et al., 2015; Libbrecht et al., 2016; Gladstad et al., 2017; Toth and Rehan, 2017), nor with the evolution of social behaviour of insects in general (Bewick et al., 2017). The role of cytosine methylation underlying gene expression differences of transient behaviours has not been assessed. More generally, it is still unknown the mechanisms underlying rapid, transient, and flexible transitions of behaviour are the same as those that are associated with longer-term behavioural transitions.

Our first goal was to identify gene expression associated with three different sources of variation; differences between individuals expressing different behaviours, differences between individuals due being with or without a mate, and differences between individuals that did or did not change their behaviour during the study (Table 1). We predicted that differences of behaviour would have a large influence on gene expression (Parker et al., 2015), followed by difference of social context (Parker et al., 2015), and the influence of individual flexibility of behaviour was largely unknown. We also predicted the possible change of expression of several pathways, including neuropeptides (Cunningham et al., 2016; 2017; Bukhari et al., 2017), neural remodelling factors (e.g, *bdnf*; Cunha et al., 2010), and genes associated with transcriptional regulation in general (Cardoso et al., 2015). Our second goal was to assess if cytosine methylation underpinned the rapid changes of gene expression seen during rapid changes of behaviour using whole genome bisulfite sequencing (WGBS) of DNA of the same males used for the gene expression experiments. Assuming cytosine methylation underpins behavioural changes, we expected to see cytosine methylation levels would change for behaviourally-responsive genes (Cardoso et al., 2015; Yan et al., 2015). We found many differences of gene expression between caring and non-caring behavioural states, fewer expression differences due to changing social contexts or individual flexibility. Very few cytosine methylation changes were associated with any of the sources of variations influencing gene expression we tested. Thus, differential expression of genes accompanies rapidly changing behaviour with a hierarchy of influences from behavioural state, social context, and individual flexibility; however, cytosine methylation does not appear to underpin any of these rapid changes and the epigenetic mechanisms that influence this process remain to be identified.

## Materials and Methods

### Parental Care of *N. vespilloides*

The parental care behaviour of *N. vespilloides* is multifaceted, easily observed, and reliably scored (Smiseth et al., 2004, 2005; Walling et al., 2008). Parental care in all burying beetle species is extensive and elaborate, including direct provisioning of regurgitated food to begging offspring (Eggert and Müller, 1997; Scott, 1998). Parental care can be uniparental or biparental, often within a species. Adults search for and bury a small vertebrate carcass on which they feed and rear offspring. Parents provide both indirect and direct care. Before young are born there is indirect care involving stripping the fur (or feathers or scales) from the carcass, forming it into a nest, and preventing microbial growth on the carcass through excretions. The latter form of indirect care also occurs after young are present, along with resource defence (Walling et al., 2008). Eggs are deposited away from the carcass while it is being manipulated into a suitable larval food resource. When eggs hatch, the larvae crawl to the carcass and reside in a small cavity excised by the parents in the carcass. Parents provide direct care by regurgitating pre-digested carrion directly to their dependent, begging offspring and by depositing enzymes into the larval cavity to provide pre-digested food for larvae in the cavity. In *N. vespilloides*, the species studied here, parental care can be provided under multiple social contexts; by both parents or either individually without influencing the survival or vigour of larvae (Parker et al., 2015). When both parents are present, females provide more direct care to offspring while males spend more time on indirect care (Smiseth et al., 2005). This system is amenable to our experimental manipulation (Table 1) as removing females while larvae are still young results in males changing to direct care (Smiseth et al., 2005). This behavioural manipulation allowed us to separate the influence of behavioural state, social context, and individual flexibility on gene expression underlying these separate forms of plasticity.

### Experimental Design and Behavioural Observations

We obtained beetles from an outbred colony of *N. vespilloides*, originating from Cornwall, UK, and maintained at the University of Georgia (Cunningham et al., 2014, 2017). This colony is augmented with new families yearly from the same origin population. We followed the protocol of Smiseth et al. (2005) to generate flexibly caring males. Unrelated female and male pairs (age 14 – 29 days) were placed into a mating box with a mouse carcass (19-21g) and 2.54 cm of moist soil. The boxes were observed every morning (approximately 9:00 am) and evening (approximately 17:00) starting at 60 h post-pairing until larvae arrived at the carcass. 21 h after larval arrival, each pair was observed using 1 minute scans for 10 minutes an hour for four observation periods. We then repeated the observation protocol 24 h later. There were two treatments on Day2: we removed half the female from pairs where the males showed no direct care on Day1 and left the pairs intact for the other half. If males were observed caring on Day 1, we left the pair intact. All pairs were observed both days regardless of treatment.

Because we were first interested in separating the influence of three factors on gene expression, we designed an experiment that manipulated males into one of four different experiences that allowed us to assess three a priori contrasts (Table 1a): The first sample, phenotypically “Flexible care”, contained males that initiated care when the female was removed. The second, “Non-flexible no-care”, contained males that never cared even if the female was removed. The third sample, “Biparental no-care”, contained males that never cared with the female present both days. The fourth sample, “Biparental care”, contained males that that always cared with the female present both days. Other samples are not available as males do not provide care on Day 2 if they do not care on Day 1 in the presence of females, and males that provide care on Day 1 rarely change to no-care on Day 2 regardless of the presence or absence of the female. To maximize power, we only selected males for analysis that showed “pure” phenotypes; that is, consistently high care or absolutely no-care throughout all observation periods.

### mRNA-sequencing (RNA-seq) Preparation, Sequencing, and Quality Control

We dissected brains from individual males as in Cunningham et al. (2014), with the exception that samples were snap frozen in liquid nitrogen after dissection and stored at -80°C. From these samples we extracted RNA and genomic DNA (gDNA) simultaneously using Qiagen’s AllPrep DNA/RNA Mini Kit (cat. # 80284; Hilden, Germany) following the manufacturer’s protocol after homogenization with a Kontes handheld pestle (Kimble Chase, Rockwood, TN, USA) to allow us to quantify both gene expression and methylation level from the same individual. We quantified RNA and gDNA with a Qubit 2.0 Flourometer (Invitrogen, Carlsbad, CA, USA) using the RNA Broad Range and dsDNA High Sensitivity protocols, respectively, following the manufacturer’s instructions.

We prepared libraries for RNA-seq with a modified Smart-seq2 protocol (Picelli et al., 2014) using a target of 80 ng of total RNA per library and barcoded with Illumina TruSeq indexes. Libraries were SPRI’ed to select for fragments between 300-1000 bp and insert size was estimated with a Fragment Analyzer Automated CE System (Advanced Analytical, Ankeny, IA, USA). We sequenced 24 samples (six from each of the four behavioural states), assigned to one of two pools to evenly distribute samples based on experimental factors across the two lanes, with a 75bp single-end (SE) protocol using to Illumina’s NextSeq500 with a High-Output flow cell targeting 35 million reads per sample at the University of Georgia’s Georgia Genomics Facility (Supplementary Table 1).

We assessed the quality of the raw sequencing reads using FastQC (v0.11.4; default settings; bioinformatics.babraham.ac.uk/projects/fastqc). We trimmed for the transposase adapter, reads based on quality (Phred > 15 at both ends), trimmed the last two base pairs of the reads due to highly skewed nucleotide frequencies, and reassessed quality of the reads using FastQC (v0.11.4) and Cutadapt (v1.9.dev1; Martin, 2011).

### Differential Gene Expression and Gene Ontology (GO) Analysis

We combined data from the four groups of males with different experiences of parental care, social context, and individual flexibility to parse the effect of different influences and refine the potential causal differences (Table 1b). We preformed three contrasts. First, we compared all those individuals that displayed parental care to those that did not, regardless of social context (Behavioural State contrast). This compares males that transition from a no-care state to a care state versus those that do not make this transition with or without the female. Next, we compared individuals in the presence of a female both days to those where a female was absent the second day, regardless of their own behaviour (Social Context contrast). This tests for the transcriptional response to difference of the social context. Finally, we tested for the transcriptional response to changes of individual behaviour regardless of social context or starting behaviour (Individual flexibility contrast). This directly compares flexible to non-flexible individuals. Taken together, then, these three contrasts lead to a description of gene expression unique to each source of variation influencing behavioural changes.

We used RSEM (v1.2.20; default settings; Li and Dewey, 2011) with BowTie2 (v2.2.9; default settings; Langmead and Salzberg, 2012) to map and quantify reads against the *N. vespilloides* Official Gene Set (OGS) v1.0 transcriptome (Cunningham et al., 2015). To better assess the completeness of the Nv OGS v1.0 before mapping, we used the updated BUSCO gene set (v2.0; default settings; Simao et al., 2015) with the Arthropoda Hidden Markov Models (2,675 HMM gene models). This gene set is defined as gene models that are present in 90% of the searched species as single-copy orthologs. We found that 2,607 (97.5%) genes were present with 2,484 (92.8%) as complete gene. Of the complete genes, 2,183 were single-copy orthologs and 301 were duplicated. A further 123 genes were fragmented.

Differential expression was estimated using both a parametric and non-parametric differential gene expression analysis to find genes that individually exhibited strong responses to our manipulation. These two methods find differential expression based on different biological signals and so can identify different sets of genes between contrasts of interest. For each analysis, we performed three contrasts (Table 1b).

We imported the expected read count per gene from RSEM into the DESeq2 package (v1.12.3; default settings; Love et al., 2014) using the tximport package (v1.0.3; Soneson et al., 2015) of R (v3.3.1; R Core Team, 2016). Following the suggested workflow of DESeq2, we preformed overall sample quality control by visual inspected for and removed two outlier samples (one flexible care, one nonflexible care) after completing quality control by visual inspection of a principal component analysis (PCA) plot using the data without regard to any factor in the study design. Statistical significance was assessed after a Benjamini-Hochberg (BH) correction of *P*-values (Benjamini and Hochberg, 1995). We used NOISeq (2.16.0; Tarazona et al., 2015) to test for differential gene expression as a non-parametric method. Following the suggested workflow of NOISeq, we preformed overall sample quality control by visual inspected for and removed one outlier sample (flexible care) after completing quality control by visual inspection of samples on PCA plot using trimmed mean of M-values (TMM) standardized data without regard to any factor in the study design, as per program guidelines. Each analysis was conducted using TMM standardized data, filtering genes with counts per million reads (CPM) <1, correcting for gene length, substituting zero gene counts with 0.01 to avoid undefined gene counts, and with 20 permutations using the NOISeqBIO function. Statistical significant was assessed after a BH correction of *P*-values. We used the union of the that were differentially expressed using DESeq2 and NOISeq genes sets for each of the three contrasts to test for enrichments of all three categories of Gene Ontology (GO) terms: biological process, molecular function, and cellular component. We used the AgriGO webserver to test for enriched GO terms (Du et al., 2010). We performed a Singular Enrichment Analysis (SEA) using Complete GO terms and a hypergeometric test with a BH correction. The complete list of GO terms assigned to all *N. vespilloides* genes was used as the background for the enrichment test.

Because genes usually act within a network, and whole networks can exhibit responses to a manipulation even if the individual genes within the network do not, we also performed a weighted gene co-expression network analysis (WGCNA). This technique also allows for the centrality of a gene to a network to be estimated with the assumption that genes deeply connected within a network are of increased overall importance because changing their expression influences many other genes. We again looked for associations with our three contrasts and the expression of gene modules between these contrasts. We used the WGCNA package of R (Storey, 2002; Langfelder and Horvath, 2008) to perform a weighted gene co-expression network analysis using default guidelines and parameters. We used the Variance Stabilized Transformation that was blind to the study design from DESeq2, with the same two outlier samples removed, as input data with genes with <10 reads in 20 samples removed, as per programs suggestion. We converted the correlation matrix of variance stabilized transformed values (DESeq2’s default transformation) to a signed adjacency matrix with an exponent of 10 and a minimum module size of 30. We tested for an association between modules and traits of interest using the biweight mid-correlation (bicor) function with a robustY setting, as per program guidelines for our data types. Modules significantly associated with traits were assessed for enrichment of GO terms as described above.

### MethylC-seq Preparation and Differential Gene- and Cytosine-Methylation Analysis

We used MethylC-seq to estimate levels of cytosine methylation associated with different behavioural states. We prepared MethylC-seq libraries following Urich et al. (2015) targeting 200 ng of gDNA as input per library. Six individuals, three each from sample group 1 & 2 (Table 1a), that we used for RNA-seq were haphazardly chosen for whole genome bisulfite sequencing. Libraries were quality controlled with the above RNA-seq protocol. We sequenced six adult samples with a 150bp single-end (SE) protocol using Illumina’s NextSeq500 with a High-Output flow cell at the University of Georgia’s Georgia Genomics Facility (Supplementary File 1).

We followed the protocol of Cunningham et al. (2015) to determine the methylation status of individual cytosines and genes that was used to survey the methylome of larval *N. vespilloides*. Briefly, we used the methylpy analysis pipeline (Schultz et al., 2015) that checks reads for adapter contamination and quality score trimming with cutadapt (v1.9dev), maps with Bowtie1 (v1.1.1; parameters: -S -k 1 -m 1 --chunkmbs 3072 --best --strata -o 4 -e 80 -l 20 -n 0), removes PCR duplicate reads with Picard (v2.4.1; default settings; broadinstitute.github.io/picard), and uses a BH corrected binomial test against the sample specific non-conversion rate of fully unmethylated lambda gDNA to call methylated cytosines. Cytosines within a region of interest (here, CDS) were aggregated and a BH corrected binomial test against the mean percentage of methylated cytosines per gene is used to call methylated genes. To estimate how conserved gene methylation status is between adult and larval life history stages, we re-analysed the six adult samples from this study and the three larvae samples from Cunningham et al. (2015; NCBI BioProject: PRJNA283826) together. To address the influence of different sequencing coverage between these samples, we restricted our analysis to genes that had at least five CpGs covered with at least three mapped reads; Cunningham et al., 2015 within the CDS regions for all nine samples (i.e., we only assessed genes with sufficient amounts of information from all samples to reduce the influence of noise from low-coverage CpGs and coverage differences between samples). A BH corrected binomial test determined the methylation status of each gene within each sample using the mean percent of methylated CpGs of all samples across all genes as the null probability. Genes identified as methylated in all adult samples and unmethylated in all larval samples were defined as adult-specific methylated genes, and vice-versa. We defined the overlap as the union of adult methylated genes compared with the union of the larval methylated genes.

We estimated differential cytosine methylation amongst the two adult behavioural states (flexible care vs. nonflexible no-care) in two different ways (qualitative and quantitative) at the gene (Patalano et al., 2015) and individual nucleotide (Libbrecht et al., 2016) levels. Our analysis was designed within an exploratory framework to capture any signal of individual cytosine or gene methylation status associated with social behaviour. For the qualitative analysis at the gene level, we assessed how many genes were consistently methylated or non-methylated in one sample group while having the opposite methylation status in other sample group. The quantitative analysis was a BH-corrected *t*-test of the proportion of methylated cytosines across a gene or a BH-corrected *t*-test of weighted methylation level across a gene (# of methylated reads/all reads mapped to a cytosine; Schultz et al., 2012) with at least 10 mapped cytosines (12,627 genes meet the minimum coverage threshold; Patalano et al., 2015).

For the qualitative analysis at the nucleotide level, we assessed how many cytosines were methylated or non-methylated in one sample group while having the opposite methylation status in the other sample group. The quantitative analysis was a BH-corrected *t*-test of the weighted methylation level (# of methylated reads/all reads mapped to a cytosine) for every cytosine that was mapped in all adult samples with at least five reads.

## Results

### Behavioural Analysis

In the sample where males were induced to shift from no-care to care (Table 1, sample 1), the percentage of observed time spent directly feeding larvae shifted from 0 (with female; Day 1) to 28.3 ± 0.4 (after female removal; Day 2). In samples where females weren’t removed but males care was observed (Table 1, sample 4), males spent 34.0 ± 5.5 % of the observation period on care in Day 1, and 35.9 ± 4.1 % of the observation period caring for larvae on Day 2. These results recapitulate those of Smiseth et al. (2005).

### Differentially Expressed Genes and Gene Co-Expression Networks

To identify differentially expressed genes and gene co-expression networks associated with changes of behaviour in different contexts, we investigated gene expression between three contrasts: behavioural state, social context, and individual flexibility (Table 1b). For the behavioural state contrast, we found 522 total differentially expressed genes using parametric analysis (Fig 1), 150 differentially expressed genes using non-parametric analysis (union of two sets is 552 genes), and seven co-expressed gene modules using WGCNA (Modules 1, 2, 5, 7, 8, 9, 10; Table 2; Supplementary File 1). For the social context contrast, we found 97 differentially expressed genes using parametric analysis, zero genes differentially expressed using non-parametric analysis, and one co-expressed gene module using WGCNA (Module 5; Table 2; Supplementary File 1). For the individual flexibility contrast, we found 17 differentially expressed genes using parametric analysis, three differentially expressed genes using non-parametric analysis (union of two sets is 19 genes), and three co-expressed gene modules using WGCNA (Modules 1, 9, 10; Table 2; Supplementary File 1). As expected, there was little overlap between the differentially expressed genes between the contrasts suggesting that we could cleanly dissect each effect (Fig 2; Supplementary File 1).

**Figure 1.**
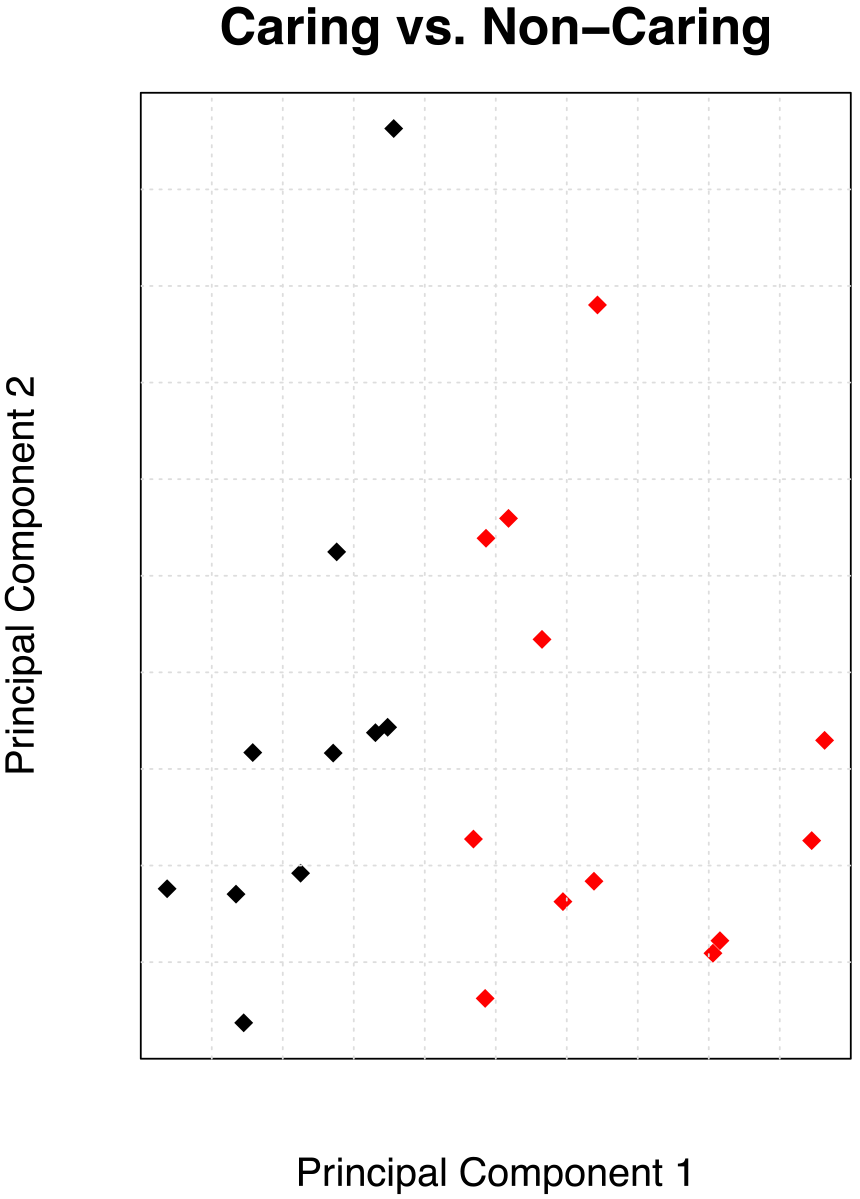
Principal component analysis of gene expression with samples coloured by Behavioural State; caring (black) vs. non-caring (red). The graph clearly shows component one as an axis of separation for this contrast.

**Table 2.** Modules of Co-Expressed Genes and their correlation with Behavioural State, Social Context, and Individual Flexibility in the context of parental care by male *N. vespilloides*.

| Module No. | No. of Genes | Behavioural State | Social Context | Individual Flexibility |
| --- | --- | --- | --- | --- |
| **0** | 1501 | -0.044 | -0.084 | 0.0013 |
| **1** | 2113 | **-0.74** | -0.085 | **-0.51** |
| **2** | 1850 | **0.48** | -0.025 | 0.13 |
| **3** | 1401 | 0.013 | -0.029 | -0.24 |
| **4** | 933 | -0.23 | -0.25 | 0.12 |
| **5** | 609 | **-0.49** | **-0.44** | -0.05 |
| **6** | 470 | 0.096 | 0.19 | 0.17 |
| **7** | 133 | **0.42** | 0.35 | 0.33 |
| **8** | 161 | **0.45** | 0.35 | 0.16 |
| **9** | 111 | **0.73** | 0.093 | **0.44** |
| **10** | 57 | **0.92** | 0.0087 | **0.51** |

Statistically significant correlations after BH-correction for multiple testing are in bold.

**Figure 2.**
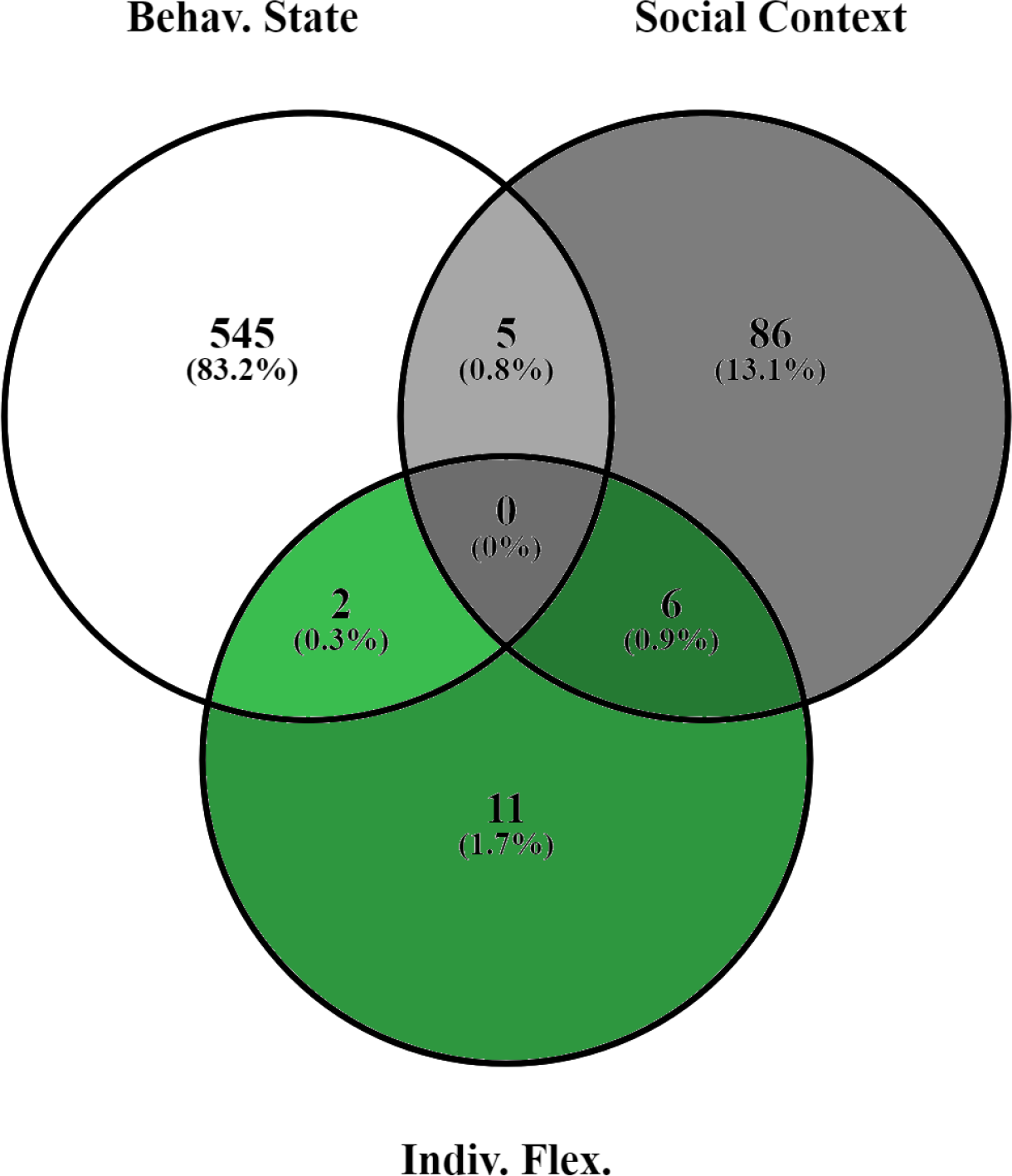
Venn diagram showing the overlap of significantly differentially expressed genes between the three contrasts analysed; Behavioural State, Social Context, Individual Flexibility.

### Functional Categories of Genes using Gene Ontology (GO) Analysis

We next used gene ontology (GO) analysis to examine the potential functions or functional categories of the genes and gene co-expression networks associated with each contrast. We found 77 GO terms enriched for the behavioural state contrast, with glutamine family amino acid metabolism, cellular aromatic compound metabolism, carboxylic acid metabolism, oxoacid metabolism, cellular amino acid biosynthetic processes, and organic acid metabolism being the most significantly associated (all *P* = 0.0063, Supplementary File 1). Only two of the seven gene co-expression networks associated with the behavioural state contrast had significant GO enrichment. Module 7 was enriched for terms related to mitochondria, cell envelope, and organelle envelope (all *P* = 0.037), whereas Module 9 was enriched for terms related to cellular amino acid metabolism, carboxylic acid metabolism, oxoacid metabolism, organic acid metabolism, and small molecule biosynthetic processes (all *P* = 0.019). Genes differentially expressed associated with variation due to difference of social context were enriched for GO terms related to only three terms; ion binding, cation binding, and metal-ion binding (all *P* = 0.011). The one gene co-expression network associated with social context had no significant GO enrichments. The differentially expressed genes of the individual flexibility contrast were not enriched for any GO terms. Of the three gene co-expression networks associated with the individual flexibility contrast, only Module 9 had enriched GO terms (see above).

### Gene and Cytosine Methylation

We investigated differences of gene or cytosine methylation to assess its relationship with flexibility in expressing care, focusing on a comparison of individuals that changed from no-care to care and those that never changed (Table 1a; sample 1 versus sample 2). This comparison should capture any mechanism associated with changes of behavioural state or individually flexibility. The genes methylated in reproductive adults overlapped highly with methylated genes in *N. vespilloides* larvae (99.4%; Fig 3). However, we found that only 2.1% of conserved adult methylated genes were also differentially expressed in any of our three contrasts (Fig 4, showing largest overlap contrast; Supplementary File 1).

**Figure 3.**
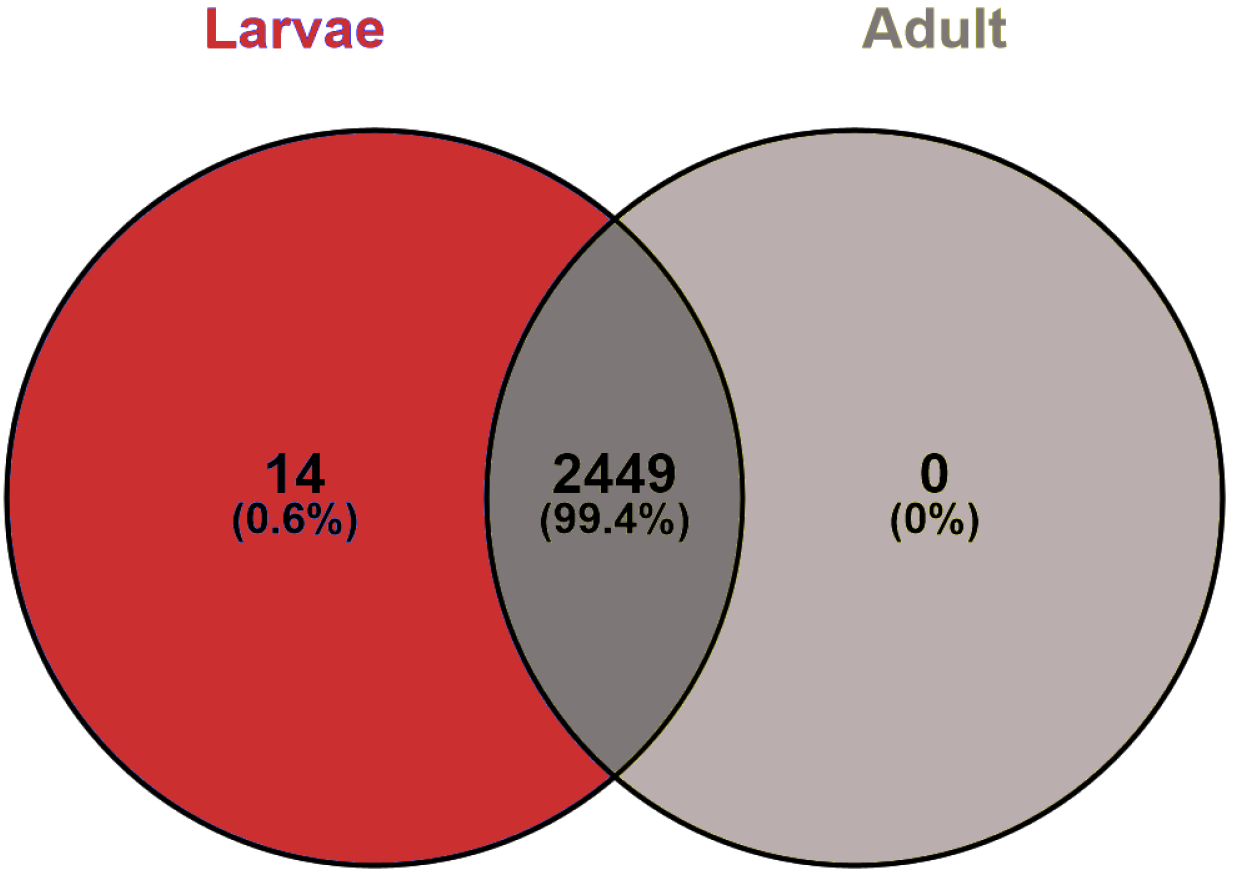
Venn diagram showing the large overlap between the methylated genes of adults and the methylated genes of larvae, using only genes that had high sequencing coverage amongst all samples to adjust for differences of sequencing depth between adult and larval samples.

**Figure 4.**
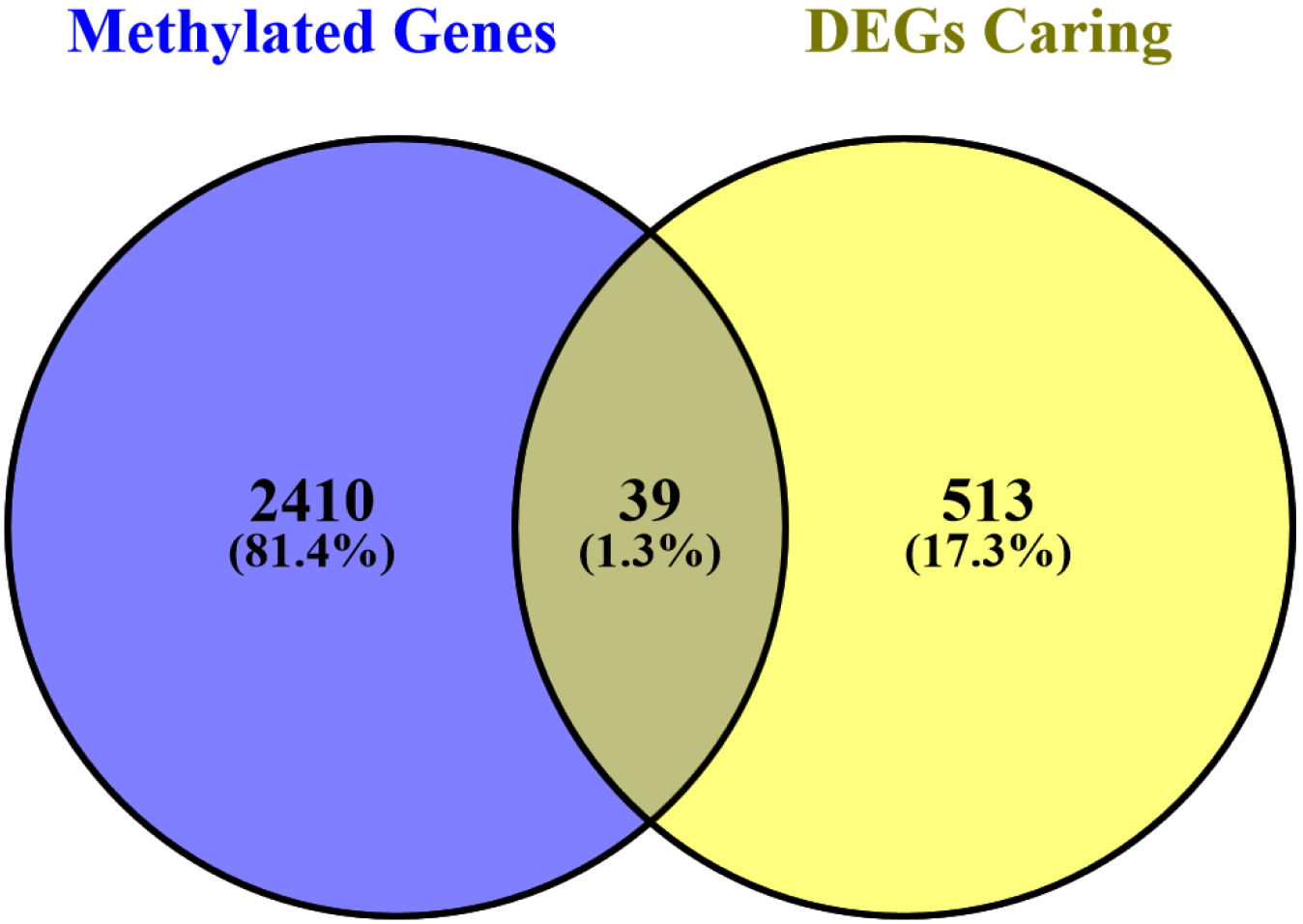
Venn diagram showing the overlap between methylated adult genes and the differentially expressed genes (DEGs) between the caring vs. non-caring contrast.

We next asked whether any methylation changes at the gene level were associated with individual flexibility of adults. We found no association between the total number of methylated genes and changes of behaviour (*t*_4_ = 0.714, *P* = 0.515). We then asked if methylation of individual genes differentiates these samples. We found 17 genes displaying a qualitative difference in methylation status. However, two methods of quantitative gene methylation analysis, percent of methylated cytosines and weighted methylation level, showed that zero and one gene, respectively, differed between flexibly expressed care and non-flexible no-care males.

It could be possible that methylation differences of individual cytosines (rather than across the entire gene body) are responsible for producing phenotypic differences. Therefore, we examined whether methylation of individual cytosines was associated with flexibly expressed care. Qualitatively, we found 460 cytosines with differing methylation status between the two groups. A permutation analysis of our samples showed that 510.5 ± 307.0 (mean ± SD) cytosines differed in methylation status. Therefore, 460 cytosines are no more than expected by chance, and provide little evidence that individual cytosine methylation is associated with behavioural state or individual flexibility. Furthermore, quantitative analysis of cytosine weighted methylation level showed only a single nucleotide (out of 56,753 methylated cytosines that had coverage in all samples) significantly associated with behavioural state or individual flexibility.

## Discussion

### Gene Expression and Differing Forms of Plasticity

Our results suggest a hierarchy of influences on gene expression during socially responsive parental care. Greater differences of gene expression were induced by manipulating behavioural states (caring vs. non-caring), fewer associated with social context, and least associated with individual variation in expressing a behaviour. The first result is consistent with the large body of studies showing differences between many behavioural states are strongly associated with gene expression differences and to a lesser extent with other factors (Zayed and Robinson, 2012; Cardoso et al., 2015; Parker et al., 2015; Toth and Rehan, 2017; Tripp et al., 2018). However, by going beyond a broad state comparison, we directly show that transcriptional architecture depends on the form of plasticity examined. The more flexible, and therefore rapid, the behavioural change the fewer gene expression changes involved.

When we assessed the functional categorization of the differentially expressed genes and gene co-expression networks, we found an abundance of metabolic related categories. Despite the abundance of GO terms related to metabolism, we do not expect these genes to reflect the energetic cost of parenting because we only sampled brains. Instead, we suggest that metabolic genes might be co-opted for a social function in *N. vespilloides*, as is argued elsewhere (Zayed and Robinson, 2012; Rittschof et al., 2014; Wu et al., 2014; Cunningham et al., 2016; Fischer and O’Connell, 2017). Alternatively, metabolic genes may be involved in neurotransmitter synthesis (Livingstone and Tempel, 1983), as many neurotransmitter pathways influence parental care (Mileva-Seitz et al., 2016). One potentially interesting candidate gene found in both the list of differentially expressed genes and as a hub gene in the gene co-expression network (Module 9) associated with caring is NK homeobox 7 (*nk7*). This gene was also one of the only genes showing evidence for positive selection in the *N. vespilloides* genome (Cunningham et al., 2015), and thus multiple lines of evidence suggest it is an important regulator of parental care behaviour. The differentially expressed genes associated with differences of social context related to ion binding, which might be associated with ion-gated channels in the brain that modulate neural activity (Simms and Zamponi, 2013). Thus, these channels may represent a candidate pathway mediating effects of the social context on behaviour. Individual flexibility of behaviour produced a clear gene expression signal associated, but the types of gene underlying this phenotype are difficult to classify. The gene co-expression network associated with flexibility is more strongly associated with caring than with individual flexibility *per se*. Individual flexibility in ants and bees is associated with morphological changes in the brain (Gronenberg et al., 1995; Groh et al., 2006), and thus we expected to detect genes annotated with neurotropic activity or neuron axon manipulation. The fact that we made no such observation suggests that gross morphological changes in the brain might only be seen in species that make permanent or developmental changes between behavioural states (Cardoso et al., 2015). It is also possible that we sampled males too late to capture the genes involved in changing gene expression, especially the immediate early genes that respond within minutes to hours to a stimulus (Cardoso et al., 2015).

### Cytosine Methylation is Not Associated with Plastic Parental Care

There is little evidence to suggest that methylation at the individual gene or individual cytosine level is associated with behavioural state or individual flexibility of male parental care of *N. vespilloides*. Adult methylated genes were highly overlapping with larval methylated genes, which indicates that gene methylation is stable across broad life history stages (and generations) encompassing widespread behavioural and physiological changes. There were few differences between at the gene or individual cytosine level between the two samples compared (Flexible Caring vs. Non-Flexible Non-Caring). Furthermore, very few (2.1%) of the adult methylated genes were also genes that were differentially expressed for any of the three contrasts of gene expression.

Our results fall in line with other studies of social insects demonstrating few differences of cytosine methylation between different behavioural states (Patalano et al., 2015; Libbrecht et al., 2016). Moreover, not all social Hymenoptera even have active DNA methylation systems (Standage et al., 2016). Cytosine methylation does not appear to be a general mechanism to regulate behavioural changes of insects (Patalano et al., 2015; Libbrecht et al., 2016; Bewick et al., 2017; Glastad et al., 2017), but it remains possible that it might regulate socially responsive gene expression of any one species. This is results is also informative because we assessed a transient behaviour, extending the range of behaviourals that cytosine methylation has been assessed to possibly influence. Furthermore, even in honey bees were many studies have reported associations between cytosine methylation and behaviour, recent research suggests that *cis*-regulatory transcription factors are strongly associated with dynamic changes of behaviour in response to social cues (Shpigler et al., 2017).

## Conclusion

Using the socially responsive and naturally variable male parental care of the subsocial beetle *Nicrophorus vespilloides*, we made a series of comparisons to understand the influence of behavioural states, social context, and individual flexibility on transcriptional architecture of a transient social behaviour. We found clear signals of gene expression after manipulating behavioural state (caring vs. non-caring), associated with social context (with or without a female mate), and to a much lesser extent with an individual’s ability to rapidly change behaviour. This suggests a complex and hierarchical influence on the transcriptional architecture of parenting behaviour by males. Research on behavioural transitions has long examined the role of single molecules, such as neuropeptides and hormones. Thus, it is perhaps no surprise that an individual’s ability to change behaviour might involve few changes of gene expression. While changes of gene expression have long been associated with changes of long-term or permanent behaviour (Zayed and Robinson, 2012; Cardoso et al., 2015), this study helps demonstrate that gene expression is also associated with rapid changes of behaviour. We find no support for an association between cytosine methylation and the expression of parental care or individual flexibility and conclude that rapid changes of cytosine methylation is not the mechanism underpinning the rapid changes of transcriptional architecture underpinning behaviour and behavioural transitions. This leads to the conclusion that, contrary to some predictions, rapid gene expression affecting behaviour may be regulated by standard processes of transcriptional control. Our work suggests that studying genetic influences underpinning changes of behavioural, perhaps one of the key attributes that defines behaviour as a unique phenotype (Bailey et al., 2018), should consider how transient the behavioural change.

## Data Availability

Data associated with this project are available at NCBI BioProject PRJNA375005. Genomic resources for *N. vespilloides* are now collated at an i5k Workspace at the National Agriculatural Library of the USDA (i5k.nal.usda.gov/nicrophorus-vespilloides).

## Acknowledgements

We thank the University of Georgia’s Georgia Advanced Computing Resource Center for computational infrastructure and technical support. Financial support for this research was provided through a National Science Foundation grant to AJM (IOS-1354358), University of Georgia’s Office of the Vice-President for Research to A.J.M. and R.J.S, and Swansea University’s College of Science to C.B.C.

## Author Contributions

CBC and AJM conceived the idea of the project. CBC, RJS, AJM designed the experiment. CBC, KMB, ECM performed the behavioural observations. CBC, ECM processed the samples. CBC, LJ performed data analysis, with assistance from RJS and AJM. CBC, KMB, AJM drafted the manuscript, which was edited by all authors.

